# AlphaFold 3-enabled *in silico* exploration of PGAM1 interactions in cancer

**DOI:** 10.1101/2025.06.23.661104

**Authors:** Jemma J. Clarke, James L. Colcombe, Daniel J. Rigden

## Abstract

Cancer cell metabolism is commonly reprogrammed to favour glycolysis over oxidative phosphorylation, even under aerobic conditions, a phenomenon known as the Warburg effect. A key enzyme implicated in this shift is phosphoglycerate mutase 1 (PGAM1), which catalyses the conversion of 3-phosphoglycerate to 2-phosphoglycerate. The human enzyme is dependent on cofactor 2,3-bisphosphoglycerate which can phosphorylate and thereby activate the enzyme at histidine 11 (H11). A recently characterised moonlighting activity of pyruvate kinase M2 (PKM2), in its monomeric or dimeric forms, leads to phosphoenolpyruvate (PEP)-dependent phosphorylation of PGAM1 at the same position. Crucially, this phosphorylation is dependent on prior tyrosine 119 (Y119) phosphorylation of PGAM1 by Src kinase, itself activated by oncogenic growth factors. Using AlphaFold 3 (AF3), this study models the conformational changes induced by PGAM1 Y119 phosphorylation and investigates the molecular basis for its role in facilitating PEP-dependent phosphorylation at H11. Structural comparisons suggest that Y119 phosphorylation induces rearrangement of the C-terminal tail of PGAM1, opening the catalytic site around H11 to enable binding of phosphoenolpyruvate (PEP). Use of molecular docking (Webina, SwissDock, and DiffDock) found that AF3 generated models of PGAM1 with Y119 phosphorylation showed enhanced binding of PEP in comparison to non-phosphorylated PGAM1. However, extensive protein–protein docking (ClusPro, AF3 multimer generation) failed to identify configurations of PGAM1 and PKM2 where catalytic sites were proximal. Overall, this study supports a model in which phosphorylation of PGAM1 at Y119 enables access to the active site for PEP, thus enhancing its enzymatic activation. These findings underscore the critical role of post-translational modifications in modulating protein function and exemplify the utility of AF3 in predicting PTM-induced structural changes relevant to cancer metabolism.

## Introduction

Cancer is frequently driven through sustained proliferative signalling and the deregulation of cellular metabolism (Hanahan, 2022). Cellular metabolism is facilitated through a plethora of post-translational modifications, a notable modification being phosphorylation (Humphrey, James and Mann, 2015). A characteristic switch in cancer cells is the production of ATP through glycolysis, rather than oxidative phosphorylation, despite the presence of oxygen. This is termed the Warburg effect, and through this cancers can break free from regular cellular regulation (Potter, Newport and Morten, 2016). The alteration of cellular signalling pathways to promote glycolysis optimises the cell for a hypoxic environment, frequently found within the tumour microenvironment. Additionally, it increases the reactive oxygen species within the cell which promotes tumour metastasis (DeBerardinis and Chandel, 2016).

Phosphoglycerate mutase 1 (PGAM1) catalyses the conversion of 3-phosphoglycerate to 2-phosphoglycerate during glycolysis, in certain cancer tissues PGAM1 is notably overexpressed (Li and Liu, 2020). Pyruvate kinase M2 (PKM2), is an enzyme that exists in monomeric, dimeric and tetrameric forms and which catalyses the conversion of phosphoenolpyruvate (PEP) to pyruvate (Wang *et al*., 2024). Tetrameric PKM2 serves to catalyse pyruvate production, whilst monomeric and dimeric forms are upregulated in highly proliferative cells such as cancers and phosphorylate PGAM1 to increase its enzymatic activity. This augmentation facilitates the Warburg effect as it increases the speed at which glucose is metabolised (Wang *et al*., 2024). Monomeric and dimeric PKM2 phosphorylates and activates PGAM1 through histidine kinase activity at H11, with the use of PEP as a phosphate donor (Wang *et al*., 2024). Wang et al. (2024) have shown that PGAM1 phosphorylation at Y119 by the Src protein is a prerequisite for PKM2 phosphorylation. Growth factors EGF, FGF and PDGF, which are proto-oncogenes, can promote Src overexpression. A summary of this interaction can be seen in Figure 1.

**Figure 1.**
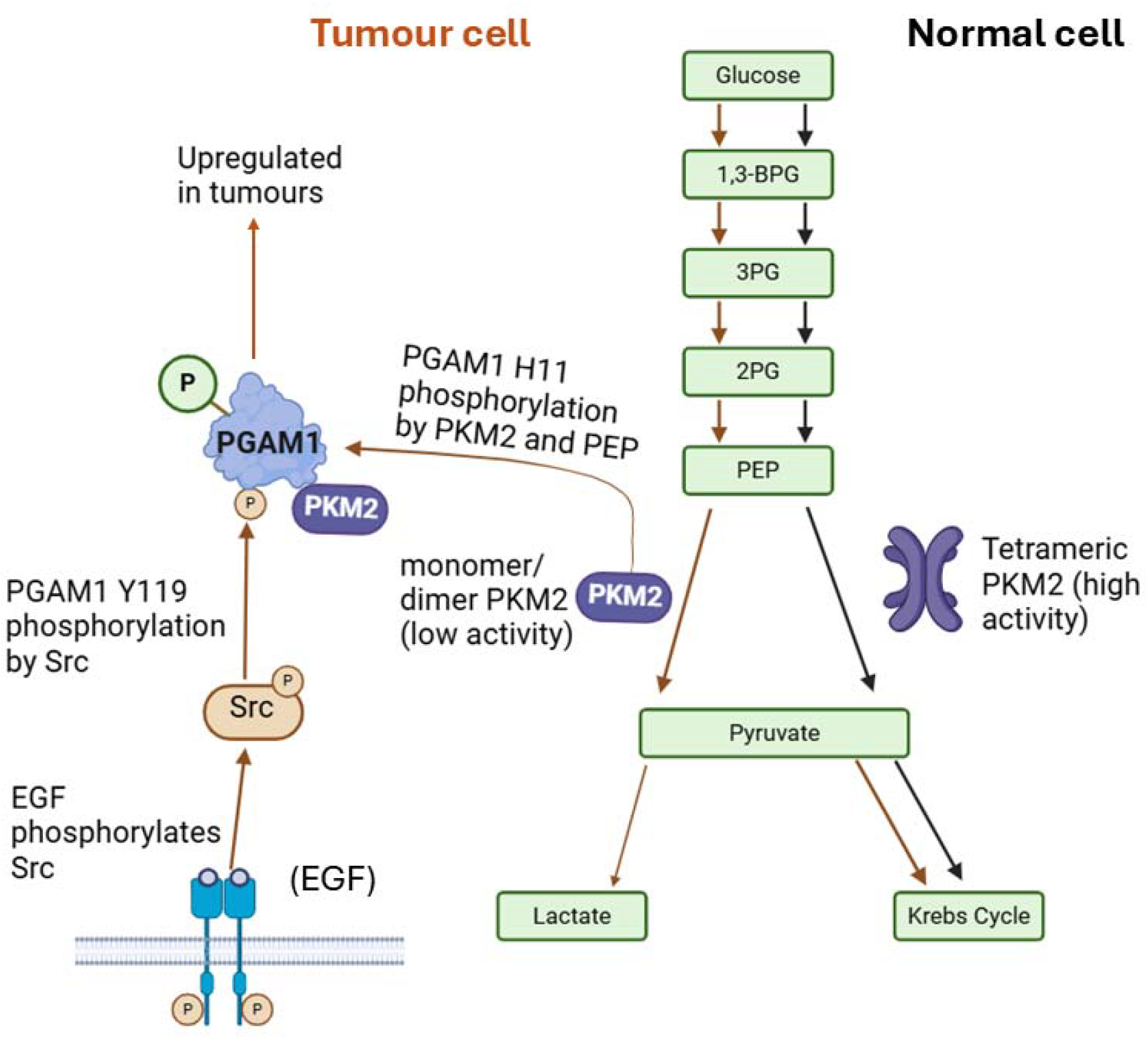
Summary of the molecular context of PGAM1 and PKM2 explored here, centred on the observations of Wang et al., (2024). PGAM1 can be phosphorylated by monomeric/dimeric PKM2 (but not the tetrameric form) which is phosphorylated itself by EGF-induced Src. The black arrow pathway shows the normal activity of tetrameric PKM2. The arrow in orange shows the pathway of monomeric/dimeric PKM2, which is phosphorylated at Y119 by active Src, and can facilitate PGAM1 phosphorylation at H11 in binding with PEP. The orange pathway is upregulated in cancer.

Here we used AlphaFold 3 and other tools to explore the molecular mechanisms potentially underlying these observations. AlphaFold 3 indeed predicted a very significant and biologically plausible conformational shift in PGAM1 upon phosphorylation of Y119, enabling docking of PEP near H11 in a fashion that was impossible in an unphosphorylated state. However, extensive modelling efforts failed to reveal how interaction of PKM2 with PGAM1 could lead to the former catalysing the phosphorylation of the latter: conceivably, greater conformational changes than could be explored bioinformatically are involved.

## Methods

### Modelling and analysis of single chains

AlphaFold3 (Abramson *et al*., 2024) was used to generate monomeric structures of phosphoglycerate mutase 1 (PGAM1) using the FASTA sequence from the Q6FHU2 entry on UniProt (The Uniprot Consortium, 2025). Two sets of five structures were generated, with or without phosphorylation at Y119.

The generated structures were compared with PyMOL. The unphosphorylated structures showed structural similarity to each other and to the 4GPI crystal structure of PGAM1 (Hitosugi *et al*., 2013). Consequently, the 4GPI model was used as a reference in this study, representing the standard dimeric PGAM1 conformation.

### Small molecule docking: docking of PEP with PGAM1 structures

To assess ligand binding, docking of phosphoenolpyruvate (PEP) with PGAM1 structures was performed using Webina (Webina: an open-source library and web app that runs AutoDock Vina entirely in the web browser)(Eberhardt *et al*., 2021). The comparison included the 4GPI crystal structure (a non-pY119 control) and the AlphaFold3-generated PGAM1 model with pY119 phosphorylation.

A 20×20×20 Å search box was defined for all docking simulations, centred on the NE2 atom of histidine 11, the residue to be phosphorylated by PEP (Rigden, 2008). The docking procedure was repeated using SwissDock (Röhrig *et al*., 2023), maintaining the same models and search box.

DiffDock (Corso *et al*., 2022) was also employed for docking simulations. Unlike Webina and SwissDock, DiffDock does not allow for specification of a search box or attraction points.

### Modelling and docking for complex structure prediction

AlphaFold3 was used to model the potential interactions between PKM2 and PGAM1. PGAM1 was modelled in its dimeric form as it is natively dimeric (Yousaf *et al*., 2023), whereas PKM2 was modelled in both monomeric and dimeric forms as these interact with PGAM1 (Wang *et al*., 2024). PyMOL was used to measure the distances between key residues.

Protein-protein docking simulations were performed using ClusPro 2.0 (Jones *et al*., 2022) to explore potential PGAM1-PKM2 interactions under different structural conditions. These conditions entail: PKM2 monomer (PDB: 3SRH) with PGAM1 dimer (PDB: 4GPI), PKM2 dimer (PDB: 6B6U), containing the S437Y mutation which promotes dimerisation, with PGAM1 dimer (4GPI), and PKM2 monomer (4QG6), which has the Y105E mutation which experimentally enhanced PGAM1 interaction (Wang et al., 2015), with the PGAM1 pY119 monomer generated using AF3. The pY119-PGAM1 monomer had its phosphate group removed in PyMOL, to maintain ClusPro 2.0 compatibility while preserving phosphorylation-induced structural changes.

In all docking simulations, attraction sites were defined to guide the progress. Specifically, residue 272 on PKM2 and residue 11 on PGAM1 were used as attraction points to enhance biologically relevant docking conformations. E272 was used as a marker of the catalytic site of PKM2, given it binds Mg^2+^ (Morgan *et al*., 2013), and H11 of PGAM1 was used as it is its active site (Hitosugi *et al*., 2013).

### Statistical analysis

Generation of box and scatter plots were performed using R (R Core Team, 2023).

## Results

### Modelling of Y119 phosphorylation of PGAM1 with AlphaFold3 produces a dramatic structural rearrangement

Superposition of the pY119 PGAM1 model with the non-phosphorylated PGAM1 model shows significant movement of the lysine tail, with the C alpha residue of Lys254 shifting by 37.2 Å (Figure 2a). Although the model confidence for the lysine tail was low (Supplementary Figure 1), this movement seems to open up the H11 site for interaction with PEP. This was found across all generated models, although the extent to which this movement was found varied. Variation in the 5 AF3 generated models was mostly found in the lysine tail, with some structural variation in the helix nearest the C-terminus (Supplementary Figure 2).

**Figure 2.**
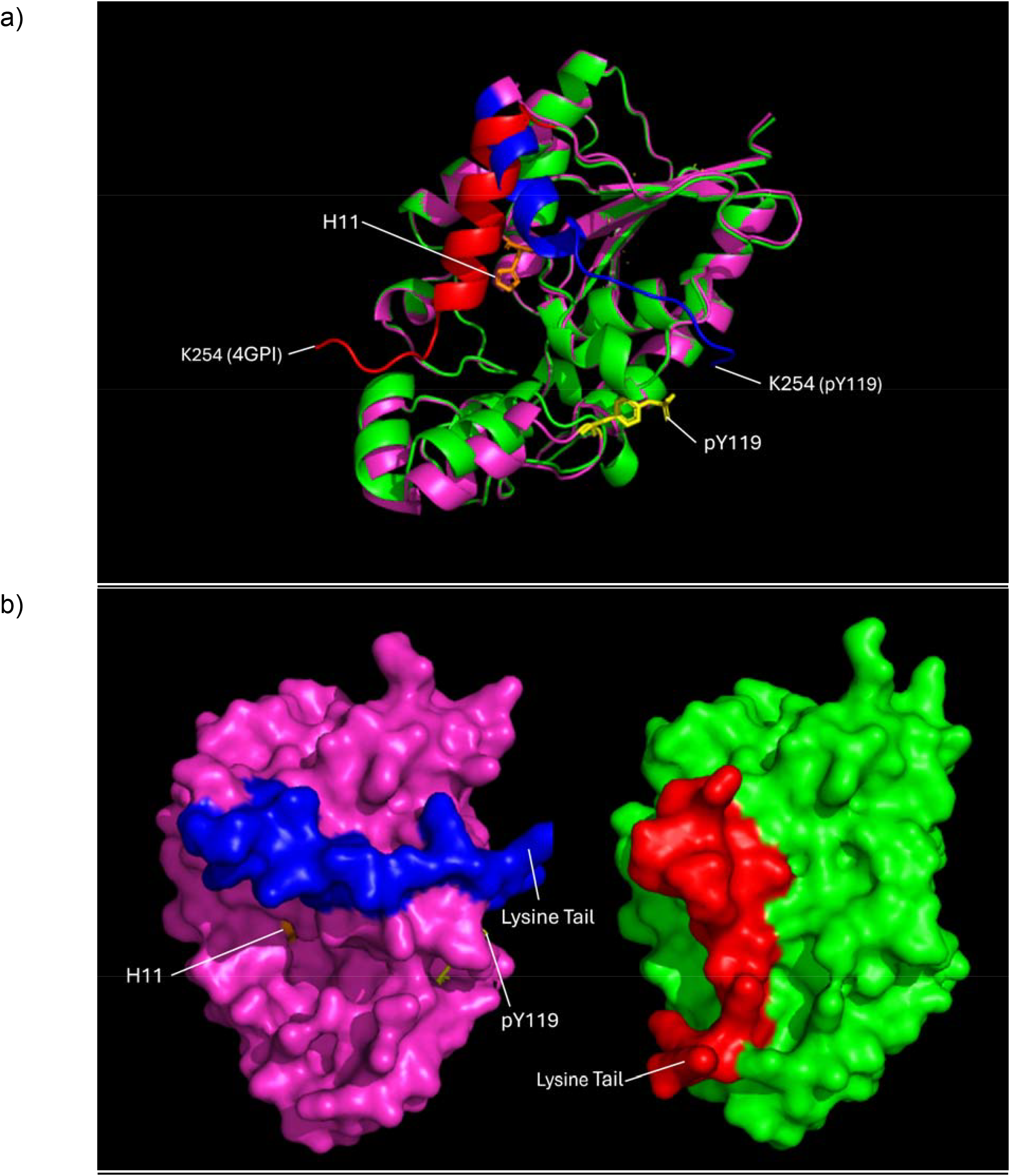
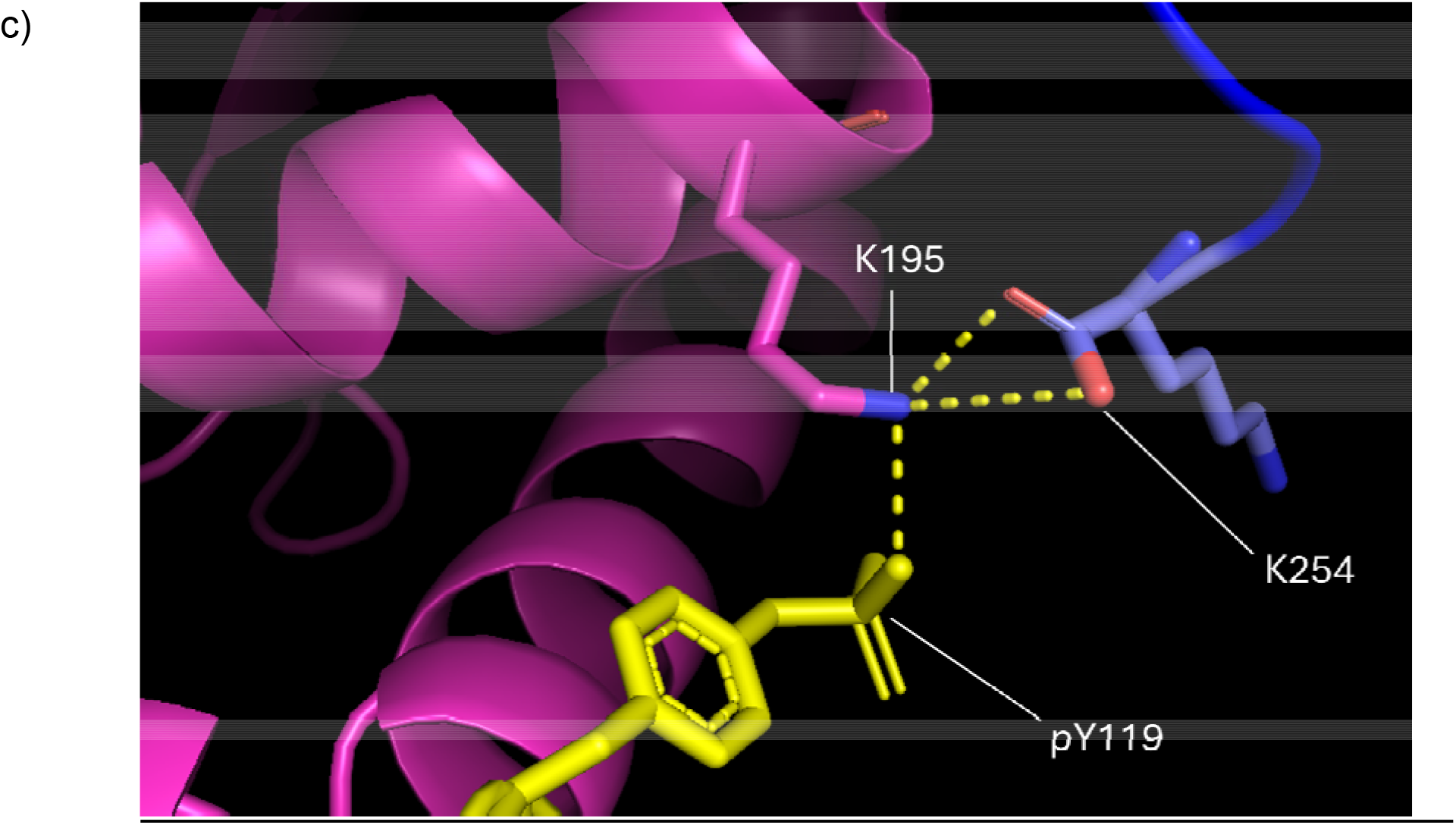
PyMOL images showing AlphaFold3 generated PGAM1 with pY119 (pink), superimposed with a non-phosphorylated PGAM1 model (4GPI). a) The pY119 residue is shown in yellow. The lysine tail of the pY119 PGAM1 model is shown in blue, contrasting with the lysine tail of the 4GPI model in red. The labelled H11 is found on both models. b) The same models are shown side by side with surface view. The left pY119 PGAM1 model has a visible cavity showing the H11 site in orange, which is covered by the lysine tail in the 4GPI model. c) A close up of the pY119 residue and its potential interaction with the lysine tail on the pY119 PGAM1 model through the intermediary side chain of K195.

Surface comparisons show that in the non-phosphorylated PGAM1 model, the lysine tail blocks the H11 site, whereas the pY119 model shows a plausible binding pocket with an open H11 residue (Figure 2b). The H11 residue had the same location in both models. Interestingly, the phosphate group of pY119 may allow for movement of the K195 side chain into the correct conformation as an intermediary to allow for interaction with the C-terminus (Figure 2c). One of the oxygen atoms of the phosphate group on Y119 was 2.8 Å away from the nitrogen of the K195 side. This nitrogen was 3.1 and 3.9 Å away from the carboxylate oxygen atoms at the C-terminus i.e. K254. Furthermore, shifting of the helices seen at the bottom of Figure 2a allows for opening of the site around the H11 residue, providing further mechanism for the enhanced binding of PEP.

### The pY119 modelled structure, but not the unphosphorylated crystal structure, can bind PEP in potentially catalytically productive modes

Observing the large structural changes induced by Y119 phosphorylation, we speculated that this could affect the binding of PEP to PGAM1. To test this, we used ligand docking simulators, including Webina, SwissDock and DiffDock, to compare PEP binding to PGAM1 with and without Y119 phosphorylation. In order to transfer its phospho group to H11, the PEP should place the phospho group in the phosphate pocket (Rigden, 2008). This was therefore used as the primary criterion to judge whether docking produced potentially catalytically productive binding.

Webina docking suggested that pY119 models allow PEP to bind closer to H11, stronger affinity, and more frequently within the phosphate binding pocket (Figure 3). SwissDock simulations furthered this idea, with the pY119 models showing more frequent instances of PEP binding appropriately compared to 4GPI (Figure 4).

**Figure 3.**
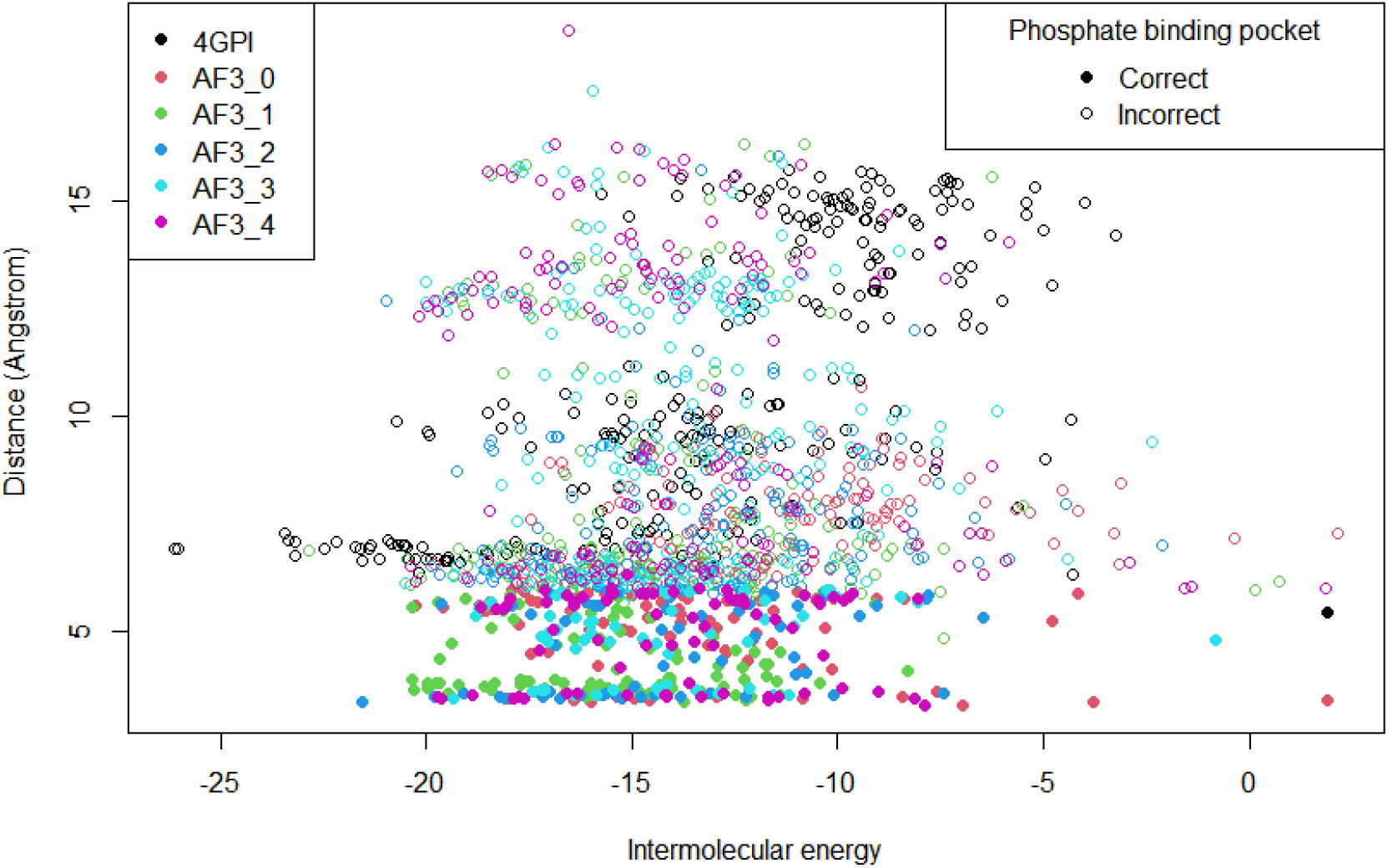
Comparison of the intermolecular energy (kJ) and distance (Å between PEP and the NE2 atom of H11 on PGAM1) of PEP binding to 5 different AF3 generated pY119 PGAM1 models and the 4GPI PGAM1 model, when docked using SwissDock.

**Figure 4.**
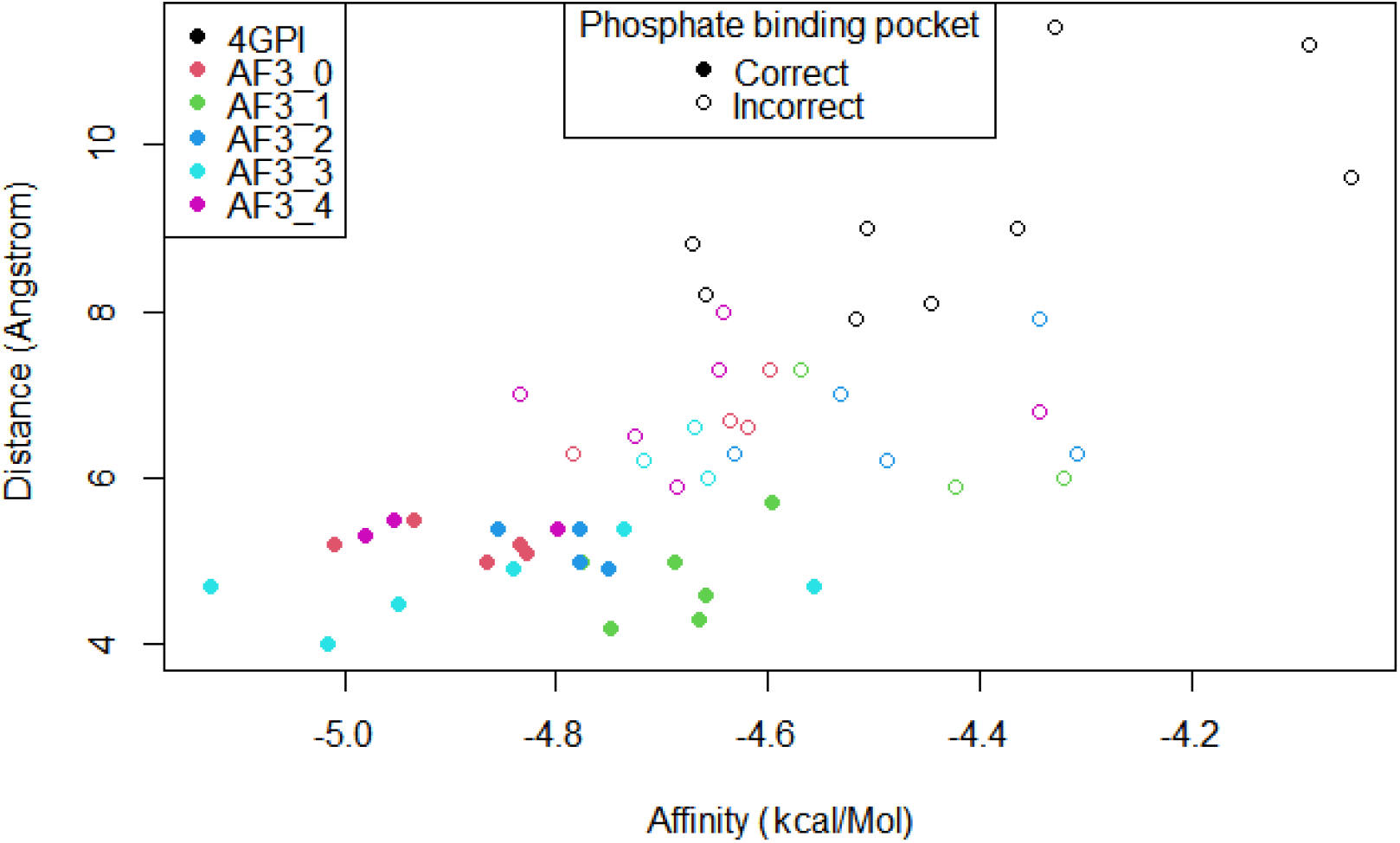
Comparison of the affinity (kcal/mol) and distance (Å between PEP and the NE2 atom of H11 on PGAM1) of PEP binding to 5 different AF3 generated pY119 PGAM1 models and the 4GPI PGAM1 model, when docked using Webina.

Conversely, DiffDock simulations showed no significant difference (Figure 5) between the pY119 and 4GPI models in the distance between PEP and the H11 site. However, it must be considered that DiffDock allowed no specification of binding or attraction points.

**Figure 5.**
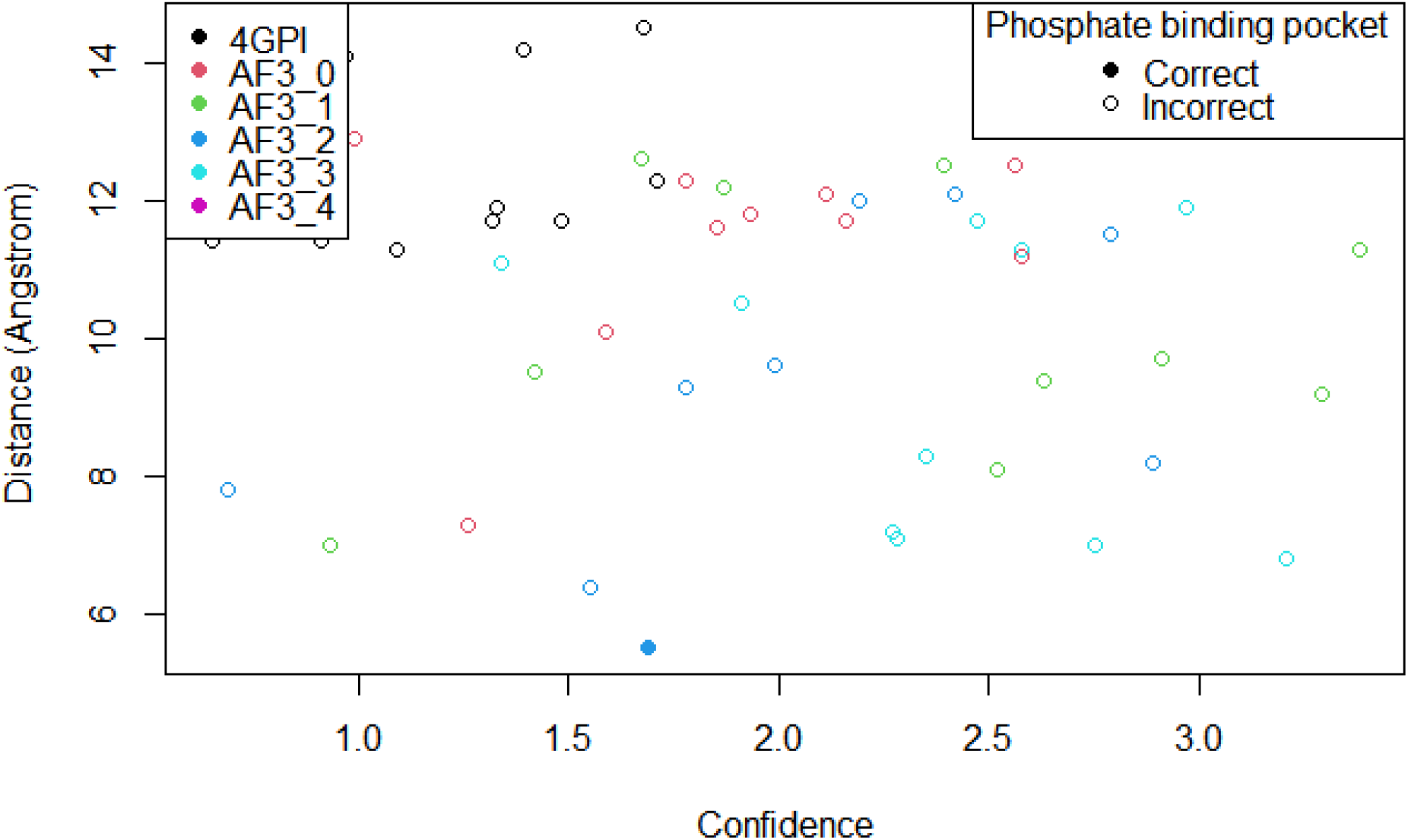
Comparison of the distance (Å) and confidence of PEP binding to 5 different AF3 generated pY119 PGAM1 models and the 4GPI PGAM1 model, when docked using DiffDock

Closest binding positions from Webina and SwissDock were found to correspond with the position of PEP that allows for interaction with the H11 residue (Figure 6), as found in previous research (Rigden, 2008). The enhanced interaction of PEP with H11 in pY119 PGAM1 models in comparison to the non-phosphorylated 4GPI model supports the proposal that pY119 phosphorylation opens up the H11 site through movement of the lysine tail and alpha helices (Figure 2).

**Figure 6.**
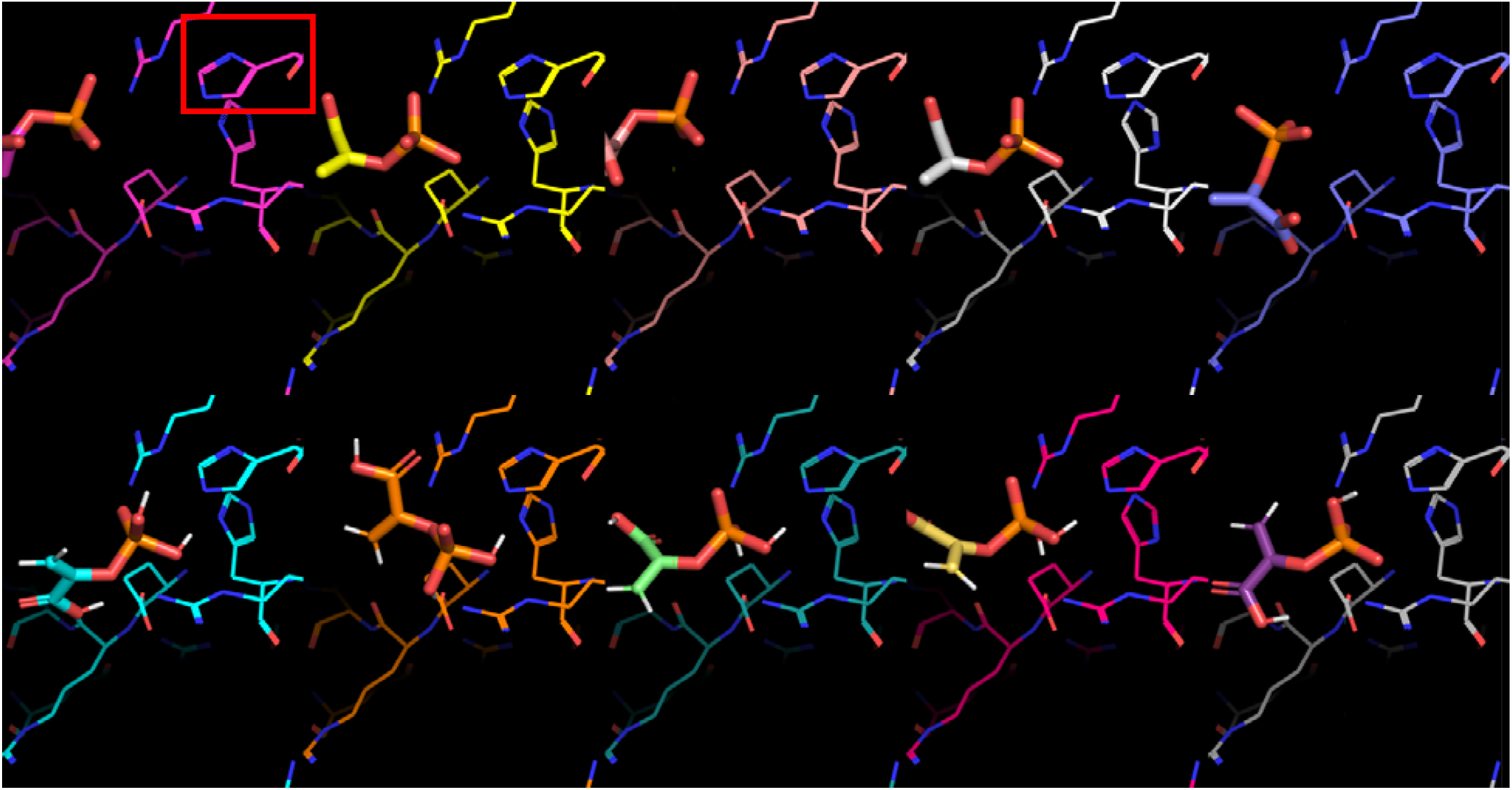
The closest positions of PEP binding to H11 (red box) in PGAM1 across pY119 AF3 models 1-5, from left to right. Top row: Docked using SwissDock. Bottom row: Docked using Webina.

### Modelling of PGAM1-PKM2 interaction fails to illuminate non-canonical phosphorylation of PGAM1

We aimed to investigate PGAM1 and PKM2 interaction to explore the hypothesis that the structures dock to allow direct transfer of the phosphate group from PEP onto PGAM1, using the catalytic machinery from PKM2. AlphaFold 3 models of PGAM1 - PKM2 interaction in various oligomers produced no complexes where catalytic sites were adjacent, thus we explored the use of ClusPro. Similarly, ClusPro docking simulations also did not show adjacent catalytic sites.

## Conclusions

The structural analysis and docking results provide a compelling possible explanation for the critical role of Src-catalyzed phosphorylation at pY119 in modulating the accessibility of the catalytic site at H11 to activation by PEP. The AF3 models of the pY119 form consistently predict a significantly more open binding site in which PEP can be placed with its phospho group in the phosphate pocket (Rigden, 2008). This contrasts with results for the unphosphorylated crystal structure, in which similarly potentially productive binding of PEP cannot occur. This study can be placed in the context of the analysis of Medvedev et al. (2025) regarding the effects of PTMs structurally on protein function. Interestingly, the conformational change predicted here, if confirmed experimentally, would be unusually large.

Extensive efforts failed to yield modes of PGAM1-PKM2 interaction which would allow for direct, PKM2-catalysed phosphorylation of PGAM1. Possibly, formation of the relevant complex requires conformational changes beyond those sampled.

To conclude, these findings collectively emphasize the importance of PTM-driven structural changes in regulating protein function and highlight the utility of advanced computational tools like AF3 in elucidating these mechanisms.

## Acknowledgment

Thanks to the University of Liverpool School of Biosciences for organising and funding the Biosciences Summer Studentship Program, and to the lab of Professor Daniel Rigden for support through this studentship.

