## Supplementary Figures for "AlphaFold 3-enabled *in silico* exploration of PGAM1 interactions in cancer"

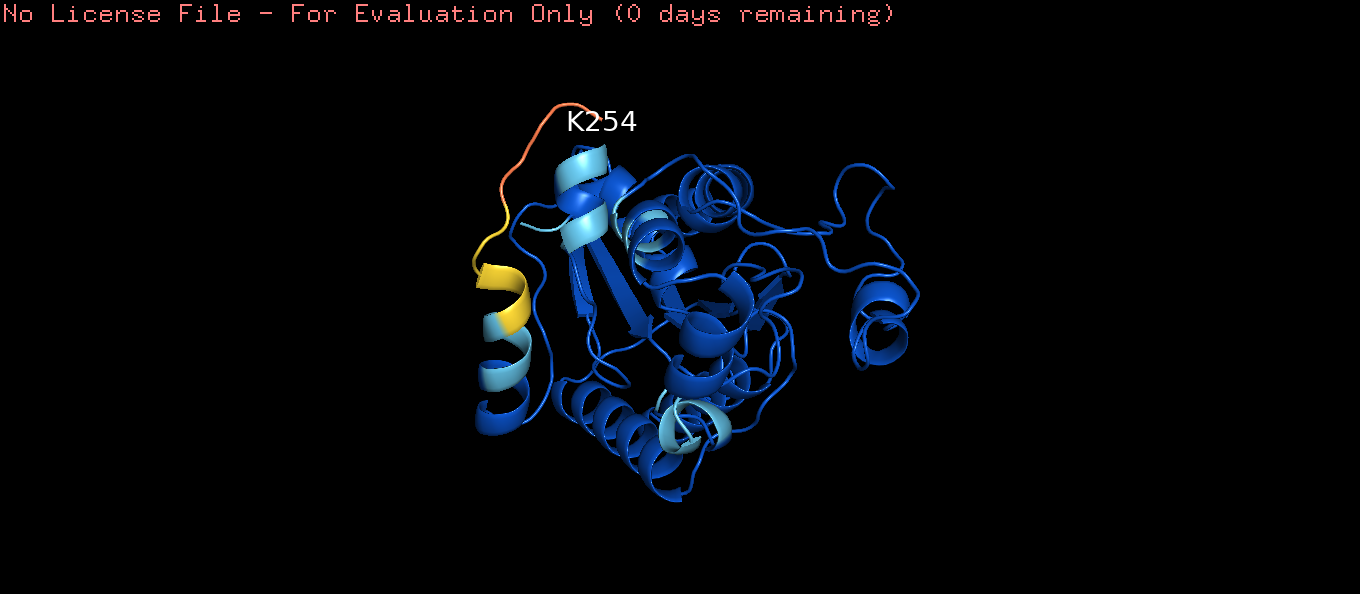

Supplementary Figure 1: PyMOL image showing AlphaFold3 generated pY119 PGAM1 coloured by plDDT value. Dark blue demonstrates very high model confidence (plDDT > 90), light blue demonstrates ‘confident’ (90 > plDDT > 70), yellow demonstrates low confidence (70 > plDDT > 50) and orange demonstrates very low confidence (plDDT < 50).

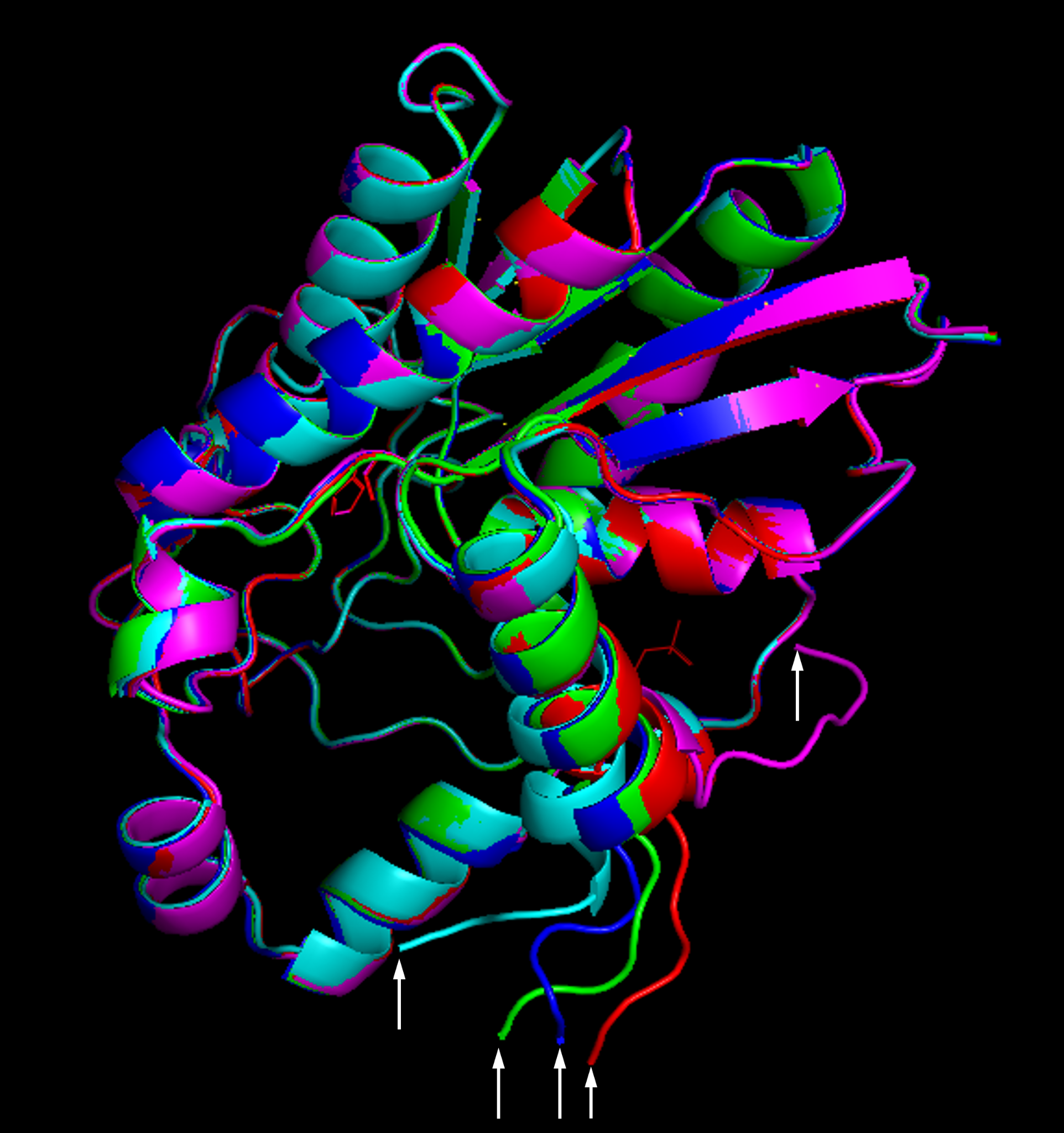

Supplementary figure 2: Superimposition of all 5 AF3 generated models of PGAM1 with Y119 phosphorylation. Arrows show K254 for each structure. Model 0 - pink, model 1 - green, model 2 - blue, model 3 - cyan, model 4 - red.
